# The EC2 domains of tetraspanins CD9, CD81, and CD151 bind to the allosteric site of integrins (site 2) and activate integrins αvβ3, α5β1 and α4β1 in a biphasic manner

**DOI:** 10.1101/2025.03.13.643145

**Authors:** Yoko K Takada, Yoshikazu Takada

## Abstract

Previous studies showed that tetraspanins activate integrins, but the mechanism of this action is unclear. We previously showed that the extracellular-2 (EC2) domains of CD9, CD81, and CD151 bind to the classical RGD-binding site (site 1) of integrin αvβ3, suggesting that they are integrin ligands. We showed that several inflammatory cytokines (e.g., CX3CL1, CXCL12, CCL5, and CD40L) bind to the allosteric site (site 2) of integrins, which is distinct from site 1, and activate integrins (allosteric activation). 25-hydroxycholesterol, a major inflammatory lipid mediator, is known to bind to site 2 and induce inflammatory signals, suggesting that site 2 plays a role in inflammatory signaling. We hypothesized that the EC2 domains activate integrins by binding to site 2. Here we describe that docking simulation predicted that CD81 EC2 binds to site 2 of αvβ3 and more strongly to site 2 of α5β1. Peptide from site 2 bound to isolated EC2 domains, suggesting that the EC2 domains bind to site 2. The EC2 domains only weakly activated αvβ3 but more efficiently activated cell surface integrins α5β1 and α4β1 on the cell surface. These results are consistent with the previous findings that these tetraspanins preferentially interact with β1 integrins. The integrin activation by the EC2 domains was increased at low EC2 concentrations and reduced as EC2 concentrations increased (biphasic), which is consistent with the findings that the EC2 domains bind to two sites (site 1 and 2). We propose that the EC2 binding to site 2 is a novel target for drug discovery.

## INTRODUCTION

Integrins are a family of cell surface αβ heterodimers and act as receptors for extracellular matrix (ECM, e.g., fibronectin and collagen), cell-surface molecules (e.g., ICAMs and VCAM1), and soluble proteins (e.g., growth factors) [1]. Integrins are known to be regulated canonically by signals from the inside of the cells (inside-out signaling) [2, 3]. Tetraspanins are a family of conserved membrane proteins that regulate cell motility, morphology, signaling, plasma membrane dynamics, and protein trafficking [4, 5]. Tetraspanins play an important role in a wide variety of physiological processes and pathogenesis of diseases. It has been proposed that tetraspanins interact with integrins and regulate integrin functions through signal transduction [6, 7]. It has been reported that expression of full-length CD81 in CD81-null U937 cells enhances α4β1 and α5β1-dependent binding to fibronectin and CD81-promoted integrin adhesiveness does not require its own ligand occupancy or ligation [8]. Also, antibodies to CD81 augment binding of erythroblasts to VCAM-1 [9]. CD9 has been identified as an activator of β1 integrins by screening shRNA library [10] and CD151 has been shown to activate α3β1 through making a tight complex with this integrin [11-14]. However, the mechanism of integrin activation by these tetraspanins has not been established.

It has been reported that the variable region of the CD151 EC2 domain (residues 186-217, helices D and E) binds to the non-ligand-binding region (residues 570-705) of the integrin α3 subunit using co-immunoprecipitation assays [12]. It has thus been proposed that tetraspanins laterally associate with integrins through non-ligand binding site of integrins [6]. We have, however, reported that the EC2 domains of CD9, CD81, and CD151 bind to the classical RGD-binding site (designated site 1) of αvβ3 [15]. This suggests that the EC2 domains are ligands for integrin αvβ3. However, how tetraspanins bind to integrins has not been fully established.

We previously reported that several cytokines such as FGF1 [16], IGF1 [17], neuregulin-1 [18], CX3CL1 [19], and secreted phospholipase A2 type IIA (sPLA2-IIA) [20] bind to site 1 of integrins. Subsequently, several cytokines induce integrin-cytokine-cognate receptor ternary comples, and this process is required for their mitogenic actions. Interestingly, we unexpectedly discovered that CX3CL1 [21], CXCL12 [22], CCL5 [23], FGF2 [24], sPLA2-IIA [25] activated integrins by direct binding to the allosteric ligand-binding site (site 2) that is distinct from the classical RGD-binding site (site 1) [21, 25]. Site 2 is on the opposite side of site 1 in the integrin headpiece [21]. Peptides from site 2 bound to CX3CL1 and sPLA2-IIA, suggesting that direct binding to site 2 is involved in the activation. We also showed that sPLA2-IIA-induced integrin activation fits well with the two-binding sites model, in which sPLA2-IIA binds to both site 1 and site 2, since activation by sPLA2-IIA is biphasic [25]. These findings suggest that the site 2-mediated integrin activation is a potential mechanism common to several pro-inflammatory proteins. The biological role of site 2 has not been fully established. It has been reported that 25-hydroxycholesterol, a major inflammatory lipid mediator, binds to site 2 and induces integrin activation and induces downstream inflammatory signals (e.g., IL-6 and TNF secretion) [26]. This suggest that site 2 is involved in inflammatory signaling. We hypothesized that the EC2 domains of CD9, CD81, and CD151 activate integrins in an allosteric manner by binding to site 2.

In the present study, we describe that docking simulation predictes that the CD81 EC2 domain binds to site 2 of αvβ3 and α5β1. Cyclic peptide from integrin site 2 bound to the CD9, CD81, and CD151 EC2 domains, suggesting that they bind to site 2. The isolated CD81 EC2 domain weakly activated αvβ3 on U937 and CHO cells. Isolated EC2 domains of CD9, CD81, and CD151 strongly activated α5β1 and α4β1 on U937 cells, suggesting that EC2 domains mediate β1 integrin activation by binding to site 2. EC2-induced integrin activation is biphasic and fits well with the idea that EC2 domains bind to site 1 and site 2 (the two site model). We propose that tetraspanin EC2 binding to site 2 may be a novel target for drug discovery in inflammation and cancer.

## RESULTS

### CD81 EC2 domain is predicted to bind to site 2 of integrin αvβ3

Previous studies showed that CD81 EC2 domain bind to the classical ligand binding site of αvβ3 (site 1) [15]. Expression of full-length CD81 activates α4β1 and α5β1 in CD81-null U937 cells [8] but the mechanism of this activation is unclear. We hypothesized that the CD81 EC2 domain bind to site 2 and allosterically activate integrins, as in the case of inflammatory cytokines (see introduction). We performed docking simulation of interaction between CD81 EC2 and integrin αvβ3 (with a closed headpiece, 1JV2.pdb). We used closed headpiece αvβ3 since the inactive conformation is well defined. The simulation predicted that CD81 EC2 binds to site 2 (docking energy -17.7 kcal/mol) (Fig. 1a). Amino acid residues involved in this interaction are shown in Table 1.

**Table 1.** Amino acid residues involved in CD81 EC2 domain-integrin αvβ3 (closed headpiece conformation) interaction (site 2).

| CD81 EC2 (1IV4.pdb, docking energy -17.5 Kcal/mol) | $\alpha V$ (1JV2.pdb) | $\beta 3$ (1JV2.pdb) |
| --- | --- | --- |
| Val123, Phe126, Tyr127, Asp137, Asp138, Asp139, Ala140, Asn141, Asn142, Lys144, Ala145, Val146, Val147, Lys148, Thr149, Phe150, Glu152, Thr153, Asp155, Ser168, Val169, Leu170, Lys171, Asn172, Asn173, Leu174, Pro176, Ser177, Gly178, Ser179, Asn180, Ile181, Ile182, Ser183, Phe186, Lys187, Lys193 | Glu15, Gly16, Tyr18, Lys42, Asn44, Gly49, Val51, Glu52, Asp83, Ser90, His91, Arg122, Arg398, Ser399, Met400 | Pro160, Val161, Ser162, Met165, Ile167, Ser168, Glu171, Glu174, Asn175, Pro186, Met187, Phe188, **Gln267, His274, Val275, Gly276, Ser277, Asp278, His280, Tyr281, Ser282, Ala283, Ser284, Thr285, Thr286, Met287** |

Amino acid residues within 0.6 nm between CD81 EC2 and $\alpha\beta 3$ were selected using Pdb viewer (version 4.1). Amino acid residues in $\beta 3$ site 2 peptide are shown in bold.

**Fig. 1.**
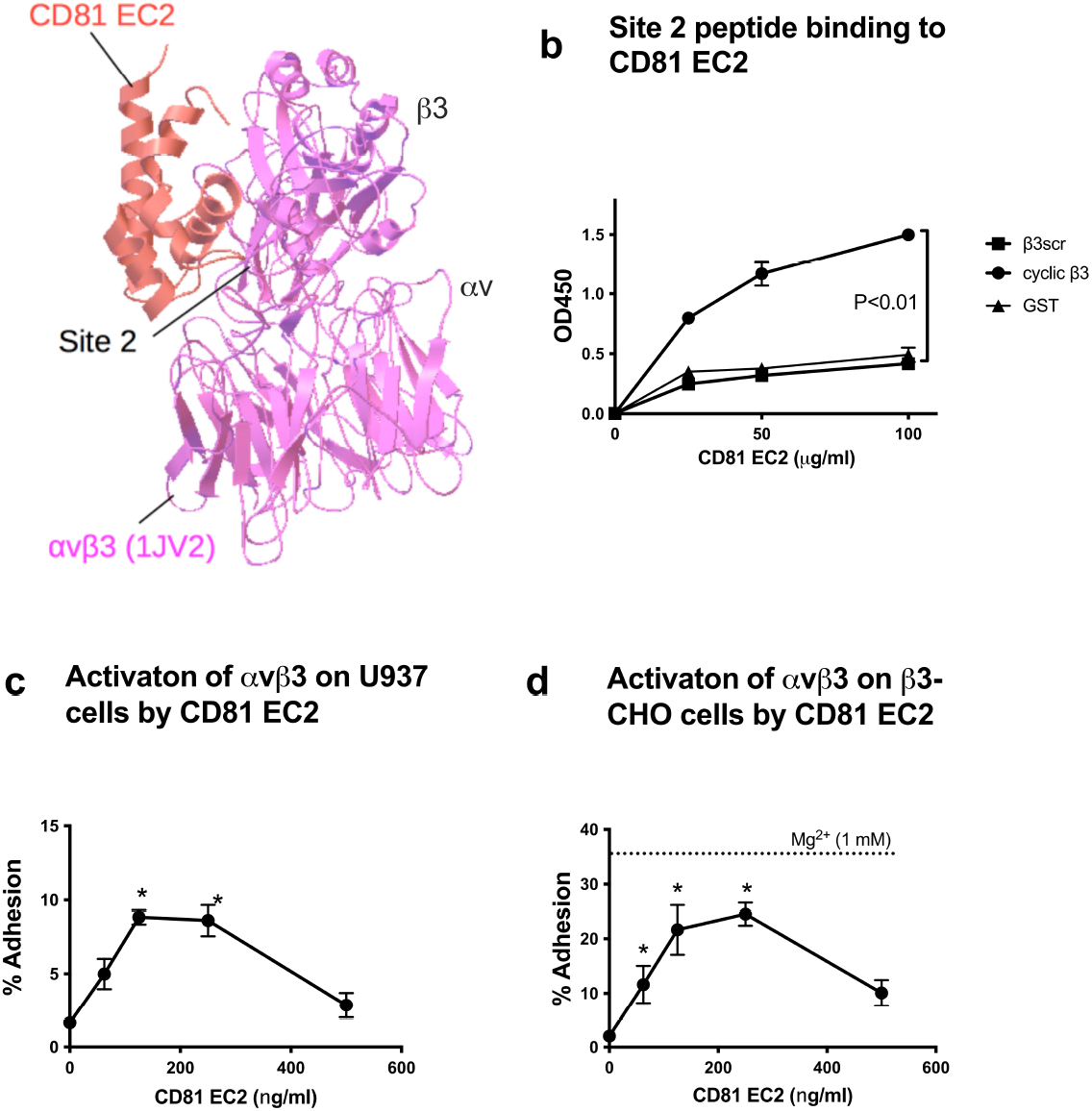
Docking simulation of interaction between CD81 EC2 and closed headpiece αvβ3. (a) Docking simulation of EC2 binding to site 2. Docking simulation was performed using the crystal structure of CD81 EC2 (PDB code 1IV5.pdb) and integrin αvβ3 (PDB code 1JV2, closed-headpiece form). A. The docking model (docking energy -17.7 kcal/mol) predicts that CD81 EC2 binds to the allosteric site (site 2) of αvβ3. (b) EC2 domains of CD81 binds to peptides from site 2 of β3. We designed disulfide-linked cyclic site 2 peptide of β3 (28-mer). Wells of 96-well microtiter plate were coated with the EC2 domains (10 μg/ml coating concentrations) and incubated with the cyclic site 2 peptide of β3 (GST fusion protein) and bound peptide was quantified using anti-GST antibodies. The values with scrambled site 2 peptide (β3scr) were subtracted. Data are shown as means +/-SEM of triplicate experiments. c and d) EC2 domain of CD81 weakly activates αvβ3 on U937 and CHO cells thar express recombinant αvβ3.

We previously reported that several pro-inflammatory proteins (CX3CL1, sPLA2-IIA, and CXCL12) bind to site 2, suggesting that they directly bind to site 2 [21, 22, 25]. Furthermore, the peptide from site 2 of β3 (21-mer) (fused to GST) directly bound to these ligands, confirming that they directly bind to site 2. We found that cyclic site 2 peptide from β3 (cyclic β3) bound to CD81 EC2 domain (Fig. 1b) better than scrambled peptide (β3scr) as a negative control, suggesting that CD81 EC2 domain specifically bind to site 2, which is consistent with the docking model.

To test if CD81 EC2 activates αvβ3, we studied if CD81 EC2 enhances binding of U937 cells (c) or CHO cells that express αvβ3 (β3-CHO) (d) to fibrinogen γ-chain fragment (γC399tr) specific ligand to αvβ3. We detected weak but significant activation.

### The CD81 EC2 domain enhances α5β1-mediated binding of U937 cells to fibronectin in a biphasic manner

Since previous studies showed that tetraspanins interact preferentially with β1 integrins, we studied if CD81 EC2 domain binds to α5β1. We performed docking simulation of interaction between CD81 EC2 and α5β1 (4wjk.pdb, with a closed headpiece) [27]). The simulation predicts that CD81 EC2 binds to site 2 of α5β1 at higher affinity (docking energy -21.95 kcal/mol) than that of αvβ3 (Fig. 2a). Nineteen out of fifty docking poses are clustered in the first cluster. Amino acid residues involved in this interaction are shown in Table 2. We studied if isolated CD81 EC2 domain activates integrin α5β1. We measured the binding of U937 monocytic cells (CD9 and CD81-deficient) to the cell-binding fragment of fibronectin (FN8-11), a specific ligand for α5β1, in the presence of the isolated CD81 EC2 domain. We confirmed that a mAb specific to α5 (KH72) effectively suppressed the binding of U937 cells to FN8-11 (Fig. 2b), suggesting that this interaction is specific to α5β1. To keep integrins inactive, we used RPMI 1640 medium, which contains high [Ca^2+^] (0.42 mM), for binding assays. As a positive control, we showed that 1 mM Mn^2+^ fully activate α5β1 in the assay medium (Fig. 2c). The CD81 EC2 (Fig. 2d) enhanced the binding of U937 cells to FN8-11 in a dose-dependent manner. Notably, at optimum concentrations CD81 EC2 enhanced α5β1 to the levels comparable to those by 1 mM Mn^2+^, suggesting that the EC2 domains likely fully activate α5β1 (Fig. 2d). Notably, the binding of U937 cells to FN8-11 was biphasic: it peaked at maximal at 15 ng/ml EC2 domains (approx. 1.5 nM) and reduced at higher concentrations (Fig. 2e). The present findings fit well with the two-sites model, in which EC2 binds to site 1 and site 2, as we reported for sPLA2-IIA-mediated integrin activation [25]. In physiological Ca^2+^ concentration in body fluid, we suspect that integrins are inactive and therefore site 1 is closed and site 2 is open. We propose that CD81 EC2 binds to site 2 first at low concentrations and activate integrins. As EC2 concentrations increase, site 2 is saturated, and EC2 domains compete with FN8-11 for binding to site 1 that is open.

**Table 2.** Predicted amino acid residues involved in CD81-α5β1 interaction (site 2)

| CD81 (1IV4, -21.95) | A5 (4wjk.pdb) site2 | $\beta 1$ (4wjk.pdb) |
| --- | --- | --- |
| Phe113, Val114, Asn115, Ile119, Asp122, Val123, Phe126, Ala130, Asn141, Asn142, Ala143, **Lys144**, Ala145, Val146, **Lys148**, Thr149, Phe150, His151, Glu152, Thr153, Leu154, Asp155, Asn172, Asn173, Leu174, Cys175, Pro176, Ser177, Gly178, Ser179, Lys187, Glu188, Lys193, Asp196, Leu197, Phe198, Ser199, Gly200, Lys201, His202 | Gly48, Leu50, Gln51, Tyr95, Ser97, Leu98, | Thr171, Val172, Met173, Ile176, Thr178, Thr179 Pro180, Ala181, Leu183, Arg184, Asn185, Gln191, Asn192, Cys193, Thr194, Thr195, Pro196, Phe197, Lys200, Val211, Glu214, Leu215, Lys218, Arg220, **His282, Glu284, Asn285, Met287, Tyr288, Thr289, Met290, Ser291, His292, Tyr293** |

Amino acid residues within 0.6 nm between CD81 EC2 and $\alpha 5\beta 1$ were selected using Pdb Viewer (version 4.1). Amino acid residues in $\beta 1$ that are in the SDL loop are underlined, and those in the $\beta 1$ site 2 peptide are shown in bold.

**Fig. 2.**
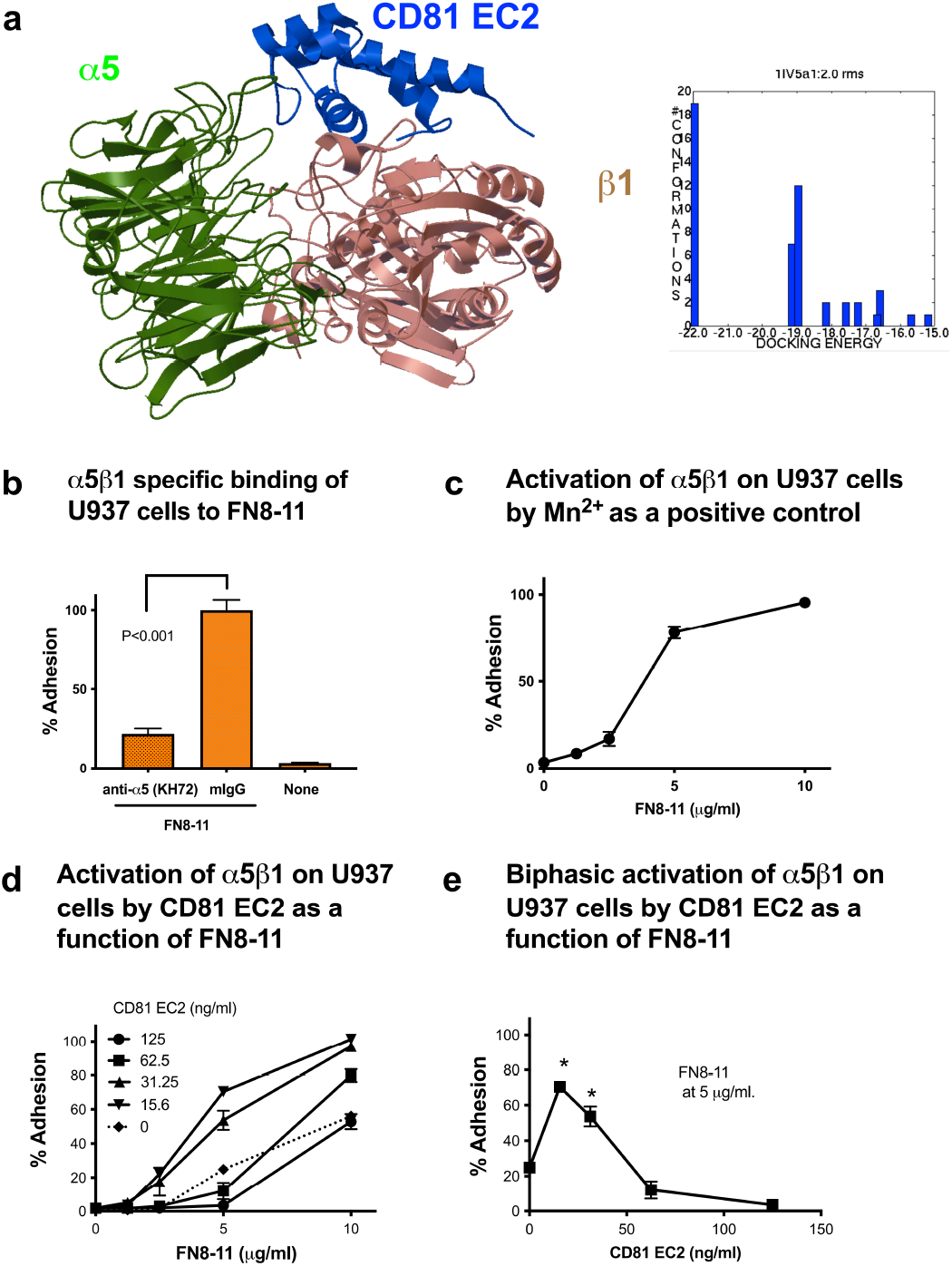
Docking simulation predicts CD81 EC2 domain binds strongly to site 2 of α5β1. (a) The docking model (docking energy -21.95 kcal/mol) predicts that CD81 EC2 binds to the site 2 of α5β1 at a high affinity. Docked poses are clustered (<0.2 RMD) and the pose in the first cluster was used. (b) Isolated EC2 domains enhanced the binding of U937 cells to the ECM ligand specific to α5β1. Wells of 96-well microtiter plate were coated with FN8-11 at 10 µg/ml coating concentrations and incubated with U937 cells in RPMI1640 medium in the presence of antibody to α5 (KH72), 1 mM Mn^2+^ (c), the EC2 domain of CD81 (d). Also, the % binding was plotted as a function of EC2 concentrations (e). Data are shown as means +/-SEM of triplicate experiments. * P<0.01 to no EC2 (e).

### The CD81 EC2 domain enhances α4β1-mediated binding of U937 cells to fibronectin in a biphasic manner

We studied if the CD81 EC2 domain activates α4β1 in a manner similar to that of α5β1. We found that binding of U937 cells to the fibronectin fragment H120, a specific ligand to α4β1 [28] was effectively suppressed by mAb specific to α4 (SG73) (Fig. 3a). This suggests that the binding of U937 to H120 is specific to α4β1. The CD81 EC2 enhanced α4β1-mediated cell binding to H120 (at 2.5 µg/ml) to the levels comparable to the activation by 1 mM Mn^2+^ (Fig. 3b), suggesting that the EC2 domains likely fully activate α4β1. The CD81 EC2 enhanced the binding of U937 cells to H120 in a biphasic manner (Fig. 3c and 3d), suggesting that CD81 EC2 allosterically activates α4β1 by binding to site 2. These findings also suggest that allosteric activation by EC2 binding to site 2 may be a common property of integrins.

**Fig. 3.**
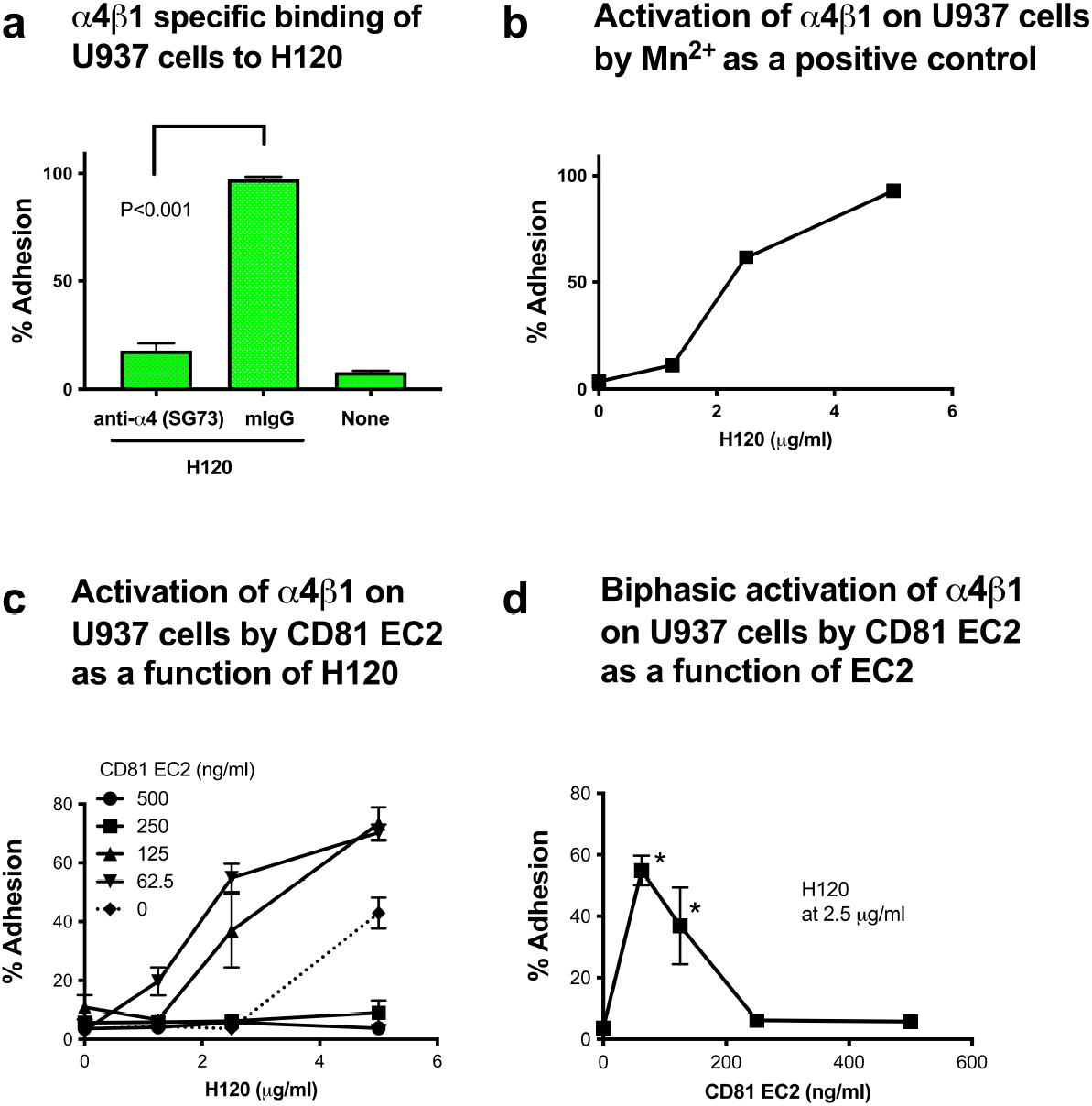
Isolated CD81 EC2 domain enhanced the binding of U937 cells to the ECM ligand specific to α4β1. Wells of 96-well microtiter plate were coated with FN H120 at 10 µg/ml coating concentrations and incubated with U937 cells in RPMI1640 medium in the presence of antibody to α4 (SG73), 1 mM Mn^2+^ (b), the EC2 domain of CD81 (c). Also, the % binding was plotted as a function of EC2 concentrations (d). Data are shown as means +/-SD of triplicate experiments. * P<0.01 to no EC2.

### The CD9 and CD151 EC2 domains enhance α4β1- and α5β1-mediated binding of U937 cells to fibronectin in a biphasic manner

The EC2 domains of CD9 and CD151 are homologous to that of CD81. We found that CD9 and CD151 EC2 bind to site 2 peptides, suggesting that they bind to site 2 (Fig. 4a and 5a). We studied if the CD9 and CD151 EC2 domains activate α4β1 and α5β1 in a manner similar to that of CD81 EC2. We found that CD9 EC2 (Fig. 4b and 4c) and CD151 EC2 (Fig. 5b and 5c) activated α5β1 in a biphasic manner. Also, CD9 EC2 (Fig. 4d and 4e) and CD151 EC2 (Fig. 5d and 5e) activated α4β1 in a biphasic manner. These findings also suggest that allosteric activation by binding to site 2 may be a common property of the EC2 domains of CD9, CD81, and CD151.

**Fig. 4.**
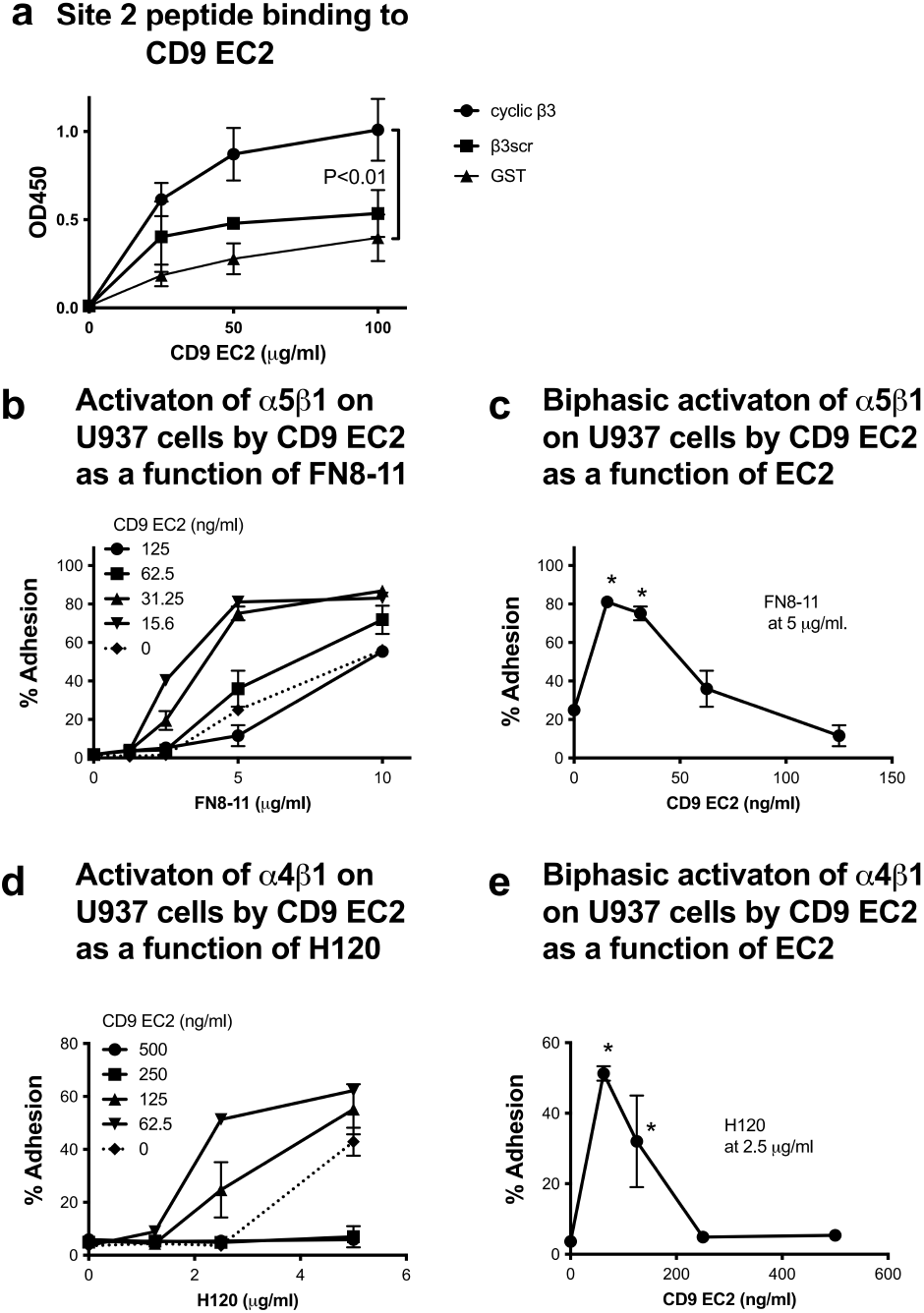
Isolated CD9 EC2 domain enhanced the binding of U937 cells to the ECM ligand specific to α5β1 and α4β1. (a) Isolated CD9 EC2 binds to site 2 peptides. (b) CD9 EC2 activates α5β1. Wells of 96-well microtiter plate were coated with FN8-11 (specific for α5β1) (b and c) H120 (specific for α4β1) (d and e) at 10 µg/ml coating concentrations and incubated with U937 cells in RPMI1640 medium in the presence of the EC2 domain of CD9. Also, the % binding was plotted as a function of EC2 concentrations (c and e). Data are shown as means +/-SD of triplicate experiments. * P<0.01 to no EC2 (c and e).

**Fig. 5.**
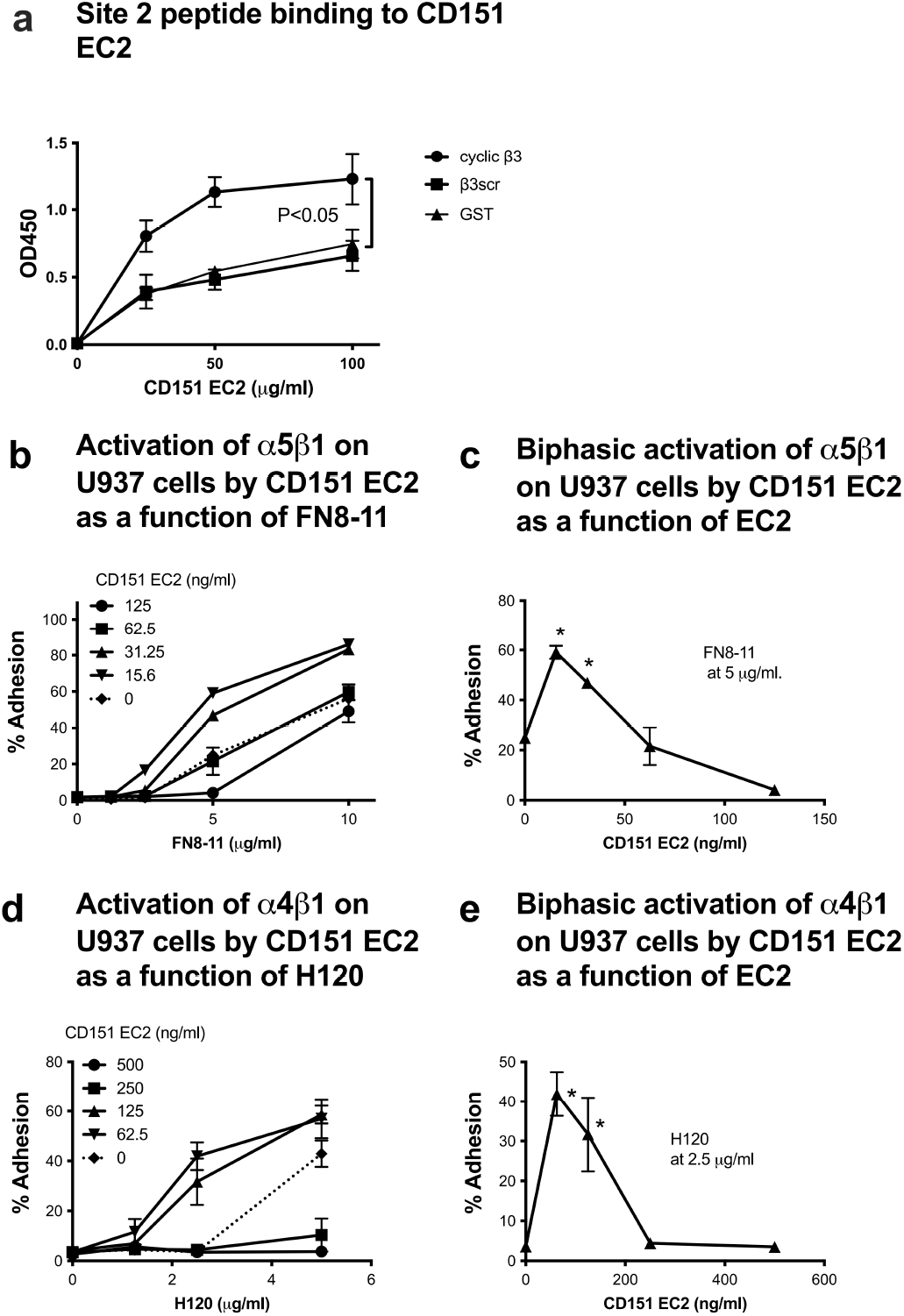
Isolated CD151 EC2 domain enhanced the binding of U937 cells to the ECM ligand specific to α5β1 and α4β1. (a) Isolated CD151 EC2 binds to site 2 peptides. (b) CD151 EC2 activates α5β1. Wells of 96-well microtiter plate were coated with FN8-11 (specific for α5β1) (b and c) H120 (specific for α4β1) (d and e) at 10 µg/ml coating concentrations and incubated with U937 cells in RPMI1640 medium in the presence of the EC2 domain of CD151. Also, the % binding was plotted as a function of EC2 concentrations (c and e). Data are shown as means +/-SD of triplicate experiments. * P<0.01 to no EC2 (c and e).

**Fig. 6.**
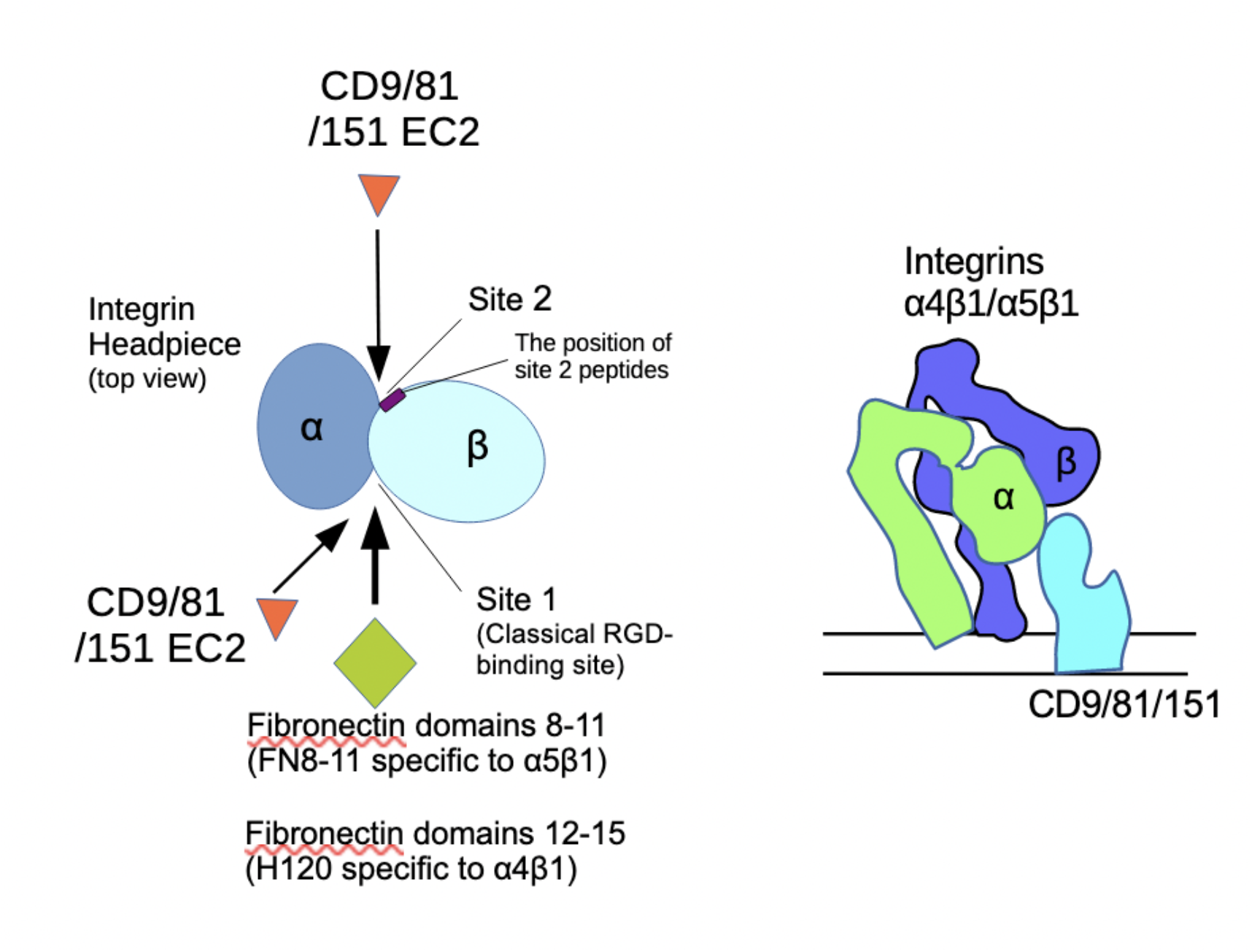
A model of integrin activation by EC2 domains. We propose a model, in which EC2 domains bind to site 2 (open in the inactive form) and activates the classical integrin ligand (RGD)-binding site (site 1) (closed in an inactive form and open in an activated form) in an allosteric manner. Since EC2 domains bind to site 1 as well, after site 2 is saturated by EC2, EC2 starts competing with ECM ligands that bind to site 1 (e.g., fibronectin or fibrinogen fragments). Thus, the effect of EC2 to integrins is biphasic.

## DISCUSSION

Previous studies showed that CD81 activate integrins [8, 29], but the mechanism of the integrin activation was unclaer. In the present study, we establish that the EC2 domains of CD9, CD81, and CD151 allosterically activate integrins α5β1 and α4β1. We recently showed that several inflammatory cytokines (e.g., FGF2, CX3CL1, CXCL12, CCL5, and CD40L) and pro-inflammatory protein sPLA2-IIA bind to site 2 and allosterically activate integrins and induce pro-inflammatory signals (see introduction). The present study suggests that the EC2 domains of CD9, CD81, and CD151 act like these inflammatory cytokines, and activate integrins. Interestingly, the EC2-induced enhancement of ligand binding was comparable to 1 mM Mn^2+^. Since EC2-mediated integrin activation was biphasic, it is expected that EC2 domains bind to site 2 at low concentrations and activate integrins and bind to site 1 at higher EC2 concentrations and compete with ECM ligands for binding to site 1 (Fig. 5). We showed that soluble EC2 domains at below 100 ng/ml (approx. 10 nM) enhanced binding through integrin α5β1 and α4β1 to the maximum levels. It appears that the EC2 domains may bind to site 2 strongly compared to sPLA2-IIA or CX3CL1 since EC2 domains required much less concentrations to activate integrins [21, 25].

Previous studies showed that CD151 is involved in tumor progression and metastasis by regulating cell motility [30] but the mechanism has not been established. We propose that allosteric integrin activation by CD151 EC2 binding to site 2 activates integrins and enhances cell migration, and at the same time enhances cytokine binding to site 1, and subsequently enhances cell migration, tumor progression, and metastasis (see Introduction). It has been reported that CD9 or CD151 silencing diminishes the relocalization of α4β1 integrin to the immunological synapse in T cell activation and reduces the accumulationof high-affinity β1 integrins at the cell–cell contact [31]. Thus, we propose the EC2 domains activate integrins through site 2 in a mechanism like that of allosteric integrin activation induced by pro-inflammatory cytokines and proteins (see Introduction).

## MATERIALS AND METHODS

### Materials

U937 cells were obtained from ATCC. b3-CHO cells have been described [32]. Hybridoma cells expressing the monoclonal antibody KH72 to a5 or SG73 to α4 were a gift from K. Miyake (University of Tokyo). GST-fusion proteins of fibronectin type III domains 8–11 (FN8-11) [19], and fibronectin H120 fragment (FN-H120) were described [21]. Fibrinogen γ-chain C-terminal domain that lacks residues 400–411 (γC399tr) was synthesized as described [33].

### Synthesis of the EC2 domains

We amplified the cDNA fragment encoding the EC2 domains of CD9 (residues 113-192), CD81 (residues 113-201), and CD151 (residues 113-221) by PCR with placenta cDNA library as a template. The amplified cDNA fragments were subcloned into the Bam HI/Eco RI site of pGEX-2T vector and PET28a vector. Synthesis of GST-EC2 fusion proteins was induced in E. coli BL21 by IPTG, and the proteins were purified from bacterial lysates using GSH affinity chromatography as described [34]. The EC2 portion was cleaved by thrombin digestion in column. The cleaved EC2 was eluted in PBS and passed through benzamidin-Sepharose to remove thrombin. 6His-tagged EC2 using PET28a vector was synthesized in E. coli BL21 by IPTG induction and purified using Ni-NTA affinity chromatography. In both cases, to remove endotoxin, affinity column was extensively washed with 1% Triton X-114 in PBS before protein elution.

### Synthesis of cyclic site 2 peptide

Cyclic site 2 peptide and scrambled peptide was synthesized as described [35]. Briefly, we introduced a disulfide linkage that connects both ends of the site 2-peptide without affecting its conformation using Disulfide by Design-2 (DbD2) software (http://cptweb.cpt.wayne.edu/DbD2/) [36]. It predicted that mutating Gly260 and Asp288 to Cys disulfide-linked cyclic site 2 peptide of β3 does not affect the conformation of the peptide. We generated C260-RLAGIVQPNDGQSHVGSDNHYSASTTMC288 29-mer β3 peptide. We found that the cyclic site 2 peptide bound to CX3CL1 and sPLA2-IIA to a similar extent to non-cyclized peptides in ELISA-type assays (data not shown).

### Binding of site 2 peptide to EC2 domains

Wells of 96-well Immulon 2 microtiter plates (Dynatech Laboratories, Chantilly, VA) were coated with 100 µl 0.1 M NaHCO3 containing EC2 for 2 h at 37°C. Remaining protein binding sites were blocked by incubating with PBS/0.1% BSA for 30 min at room temperature. After washing with PBS, GST-tagged site 2 peptides were added to the wells and incubated in PBS for 2 h at room temperature. After unbound GST-tagged site 2 peptides were removed by rinsing the wells with PBS, bound GST-tagged site 2 peptides were measured using HRP-conjugated anti-GST antibody and peroxidase substrates.

### Integrin activation by EC2 domains

Binding assays were performed as described previously [19]. Briefly, wells of 96-well Immulon 2 microtiter plates were coated with 100 µl PBS containing FN8-11, H120, γC399tr and were incubated for 1 h at 37°C. Remaining protein binding sites were blocked by incubating with PBS/0.1% BSA for 30 min at room temperature. After washing with PBS, cells in 100 µl RPMI 1640 medium were added to the wells and incubated at 37°C for 1 h. After unbound cells were removed by rinsing the wells with the medium used for binding assays, bound cells were quantified by measuring endogenous phosphatase activity [37]. Cells were preincubated with the EC2 domains for 10 min at room temperature before adding to wells. When integrins were activated by Mn^2+^, we used Tyrode-Hepes buffer plus 1 mM Mn^2+^ instead of PRMI 1640.

### Docking simulation

Docking simulation of interaction between CD81 EC2 (PDB code 1IV5.pdb) and integrin αvβ3 (PDB code 1JV2, closed-headpiece form) or α5β1 (4wjk.pdb) was performed using AutoDock 3.05 as described [16]. Cations were not present in αvβ3 during docking simulation [16, 20].

### Other methods

Treatment differences were tested using analysis of variance and a Tukey multiple comparison test to control the global type I error using Prism 10 (Graphpad Software).

## Acknowledgements

This project was supported by NIH R33CA196445. The contents reported/presented within do not represent the views of the Department of Veterans Affairs or the United States Government.

## Author Contributions

YKT performed the experiments. YT conceived the experiments and performed docking simulation and wrote the manuscript.

